# Plasticity of olfactory bulb inputs mediated by dendritic NMDA-spikes in piriform cortex

**DOI:** 10.1101/2021.05.25.445578

**Authors:** Amit Kumar, Edi Barkai, Jackie Schiller

**Affiliations:** Department of neuroscience, Technion Medical School, Bat-Galim, Haifa 31096, Israel; Sagol department of Neurobiology, Faculty of Natural Sciences, University of Haifa, Haifa, Israel

## Abstract

The piriform cortex (PCx) is essential for learning of odor information. The current view postulates odor learning in the PCx is mainly due to plasticity in intracortical (IC) synapses, while odor information from the olfactory bulb carried via the lateral olfactory tract (LOT) is “hardwired”. Here we revisit this notion by studying location and pathway dependent plasticity rules. We find that in contrast to the prevailing view, synaptic and optogenetically activated LOT synapses undergo strong and robust long-term potentiation (LTP) mediated by only few local NMDA-spikes delivered at theta frequency, while global spike timing dependent plasticity protocols (STDP) failed to induce LTP in these distal synapses. An inverse result was observed for more proximal apical IC synapses; they undergo plasticity with STDP but are refractive to local NMDA-spike protocols. These results are consistent with a self-potentiating mechanism of odor information via NMDA-spikes which can form branch-specific memory traces of odors.

## Introduction

The piriform cortex (PCx) is a main cortical station in olfactory processing, receiving direct odor information from the olfactory bulb via the lateral olfactory tract (LOT) as well as higher brain regions and is thought to be important for odor discrimination and recognition. Pyramidal neurons in the PCx serve as the main integration units within which the discrete molecular information channels are hypothesized to be recombined to form an “odor object”. PCx pyramidal neurons receive two spatially distinct inputs which terminate on different compartments of their apical dendrites: first, the direct afferent excitatory inputs from olfactory bulb (OB) via the LOT, which terminate mainly on distal apical dendrites (layer Ia). Second, intracortical excitatory inputs (IC) from local PCx neurons and other cortical areas, which target the more proximal portions of the apical dendritic tree (layers Ib and II) and basal dendrites (Bekkers and Suzuki, 2013; Haberly, 1985; Haberly, 2001; Haberly and Price, 1977; Isaacson, 2010; Neville and Haberly, 2004; Suzuki and Bekkers, 2006).

Synaptic plasticity rules have crucial role in determining the way cortical networks acquire, organize and store information. It is postulated that PCx mediates learning and recall of olfactory information (Franks and Isaacson, 2005; Haberly, 1985; Haberly, 2001; Saar et al., 1998, 2001). Previous studies have shown that similar to neocortical and hippocampal pyramidal neurons (Bliss and Collingridge, 1993; Feldman, 2012; Sjostrom et al., 2007), IC synapses of PCx pyramidal neurons undergo plasticity changes. NMDA-R dependent long-term potentiation (LTP) was robustly observed in IC synapses both when stimulated by theta rhythm alone (Haberly et al., 1994), or when paired with either a burst of LOT activation (Kanter and Haberly, 1990, 1993; Kanter et al., 1996), or back-propagating action potentials using spike timing dependent protocols (STDP) (Johenning et al., 2009). In line with the importance of IC plasticity changes in learning, it was shown that following olfactory rule learning in-vivo, PCx pyramidal neurons exhibited increase in IC synaptic strength and excitability (Lebel et al., 2001; Saar and Barkai, 2003, 2009; Saar et al., 2001).

In contrast to IC inputs, plasticity of afferent LOT synapses is yet an unresolved issue (Haberly et al., 1994; Jung et al., 1990; Roman et al., 1993). NMDA dependent LTP using theta burst stimulation of LOT inputs was less robust and weak. Typically, 10-15% potentiation was observed in afferent fibers after a single theta burst train, with saturation of 12-25% after multiple trains (Franks and Isaacson, 2005; Jung et al., 1990; Kanter and Haberly, 1990; Poo and Isaacson, 2007; Roman et al., 1993). STDP protocols failed altogether to induce LTP in LOT synapses because of the severe attenuation of back propagating action potentials (BAPs) to distal apical dendrites of PCx pyramidal neurons, where LOT inputs terminate (Haberly, 2001; Neville and Haberly, 2004). Currently, LOT synapses located in distal apical dendrites are considered for the most part to be “hardwired” after the sensory critical period (Bekkers and Suzuki, 2013; Franks and Isaacson, 2005), and robust NMDA-R dependent LTP is limited to a brief postnatal time window of approximately 4 weeks (Franks and Isaacson, 2005; Poo and Isaacson, 2007), which is in contrast to IC synapses that retain the capacity for plasticity throughout adulthood. Yet, the notion of LOT synapse stability in adulthood is in conflict with reports obtained in-vivo during complex olfactory learning (Cohen et al., 2008; Cohen et al., 2015; Patneau and Stripling, 1992).

We have recently reported the generation of local NMDA-spikes in distal apical dendrites by activation of LOT synapses (Kumar et al., 2018). These dendritic NMDA-spikes generate large localized calcium transients, which can serve as post synaptic signals for LTP induction in the activated LOT synapses. In support of this hypothesis, it was recently shown that NMDA-spikes were critical for inducing LTP in the dendrites of CA3 pyramidal neurons (Brandalise et al., 2016) and in layer 2-3 pyramidal neurons of somatosensory cortex in-vivo (Gambino et al., 2014; Gordon et al., 2006). We thus hypothesized that synchronized firing of spatially clustered afferent LOT synapses, can generate local NMDA-spikes, that in turn can trigger LTP of the activated synapses in distal dendrites of PCx pyramidal neurons.

Consistent with this hypothesis and settling the inconsistencies in the literature, we found local NMDA-spikes generated in distal apical dendrites delivered at 4 Hz mimicking the exploratory sniff cycle in rats (Wilson, 1998), can induce robust and strong LTP (>200 %) of LOT inputs impinging on distal apical dendrites of PCx pyramidal neurons. Only few NMDA-spikes (as low as 2 NMDA-spikes) were required for potentiation of LOT inputs. As previously described, STDP protocol failed to induce LTP of LOT synapses (Johenning et al., 2009). We have observed a mirror image for proximal apical IC synapses. Potentiation mediated by NMDA-spikes, was specific to the LOT synapses and was not effective for potentiating IC inputs innervating more proximal regions of the apical dendritic tree. In accordance with previous reports (Johenning et al., 2009), STDP was efficient at potentiating these synapses.

## Results

The inputs to PCx are highly segregated along the axis of apical dendrites in principle pyramidal neurons. This raises the possibility of location specific plasticity rules as was previously shown for neocortical pyramidal dendrites (Gordon et al., 2006; Kampa et al., 2007; Letzkus et al., 2006, 2007; Lisman and Spruston, 2010; Sandler et al., 2016; Sjostrom and Hausser, 2006; Sjostrom et al., 2008). In this work we tested both global spike timing dependent plasticity protocols as well as local induction protocols mediated by dendritic NMDA-spikes (Kumar et al., 2018).

### Location dependent STDP in apical and basal dendrites of PCx pyramidal neurons

To directly test the location dependent capability of PCx dendrites to undergo LTP with an STDP protocol, we performed whole-cell patch-clamp voltage recordings from the soma of layer II pyramidal neurons as determined from the Dodt contrast image and somatic firing pattern (Suzuki and Bekkers, 2006). Neurons were loaded with calcium sensitive dye OGB-1 (200 µM) and CF633 (200 µM) to enable visualization of dendrites using a confocal microscope. We used focal synaptic stimulation to activate synaptic inputs in distal LOT (299 ± 6.29 µm from soma) synapses located in layer Ia (Figure 1A) or more proximally (139.33 ± 5.15 µm from soma) in layer Ib (Figure 1E) to activate predominantly the IC synapses (Bekkers and Suzuki, 2013; Isaacson, 2010; Kumar et al., 2018; Suzuki and Bekkers, 2011). LTP was induced by pairing a single EPSP (1.88 ± 0.23 mV) with a burst of postsynaptic BAPs at 20 Hz (a sequence of three BAPs @ 150 Hz repeated 3 times at 20 Hz; Each sequence was repeated 40 times at 5 seconds interval) which simulates the beta oscillation frequency of odor-evoked synaptic activity in PCx (Johenning et al., 2009; Poo and Isaacson, 2007). (Figure 1B). In accordance with a previous report (Johenning et al., 2009), this paradigm failed to induce LTP in distally activated apical synapses at layer Ia where most of the LOT inputs terminate (Figure 1C, D; the EPSP amplitude after the induction protocol was 103.06 ± 2.81 % of the control EPSP; p = 0.7658; n = 11). Next, we replaced APs evoked by somatic current injection with postsynaptic APs evoked by synaptic stimulation in layer IIa (Figure supplement 1). When paired with an EPSP (as described in Figure 1B), this STDP protocol also failed to potentiate distally located layer Ia inputs (104.99 ± 3.86% of control; p = 0.7577, n=3). In sharp contrast, using the same protocol but activating IC instead of LOT synapses at more proximal apical dendritic locations induced robust potentiation of layer Ib inputs, (Figure 1F and 1G; 168.93 ± 6.54% of control EPSP; p = 0.01387, n = 6).

**Figure 1:**
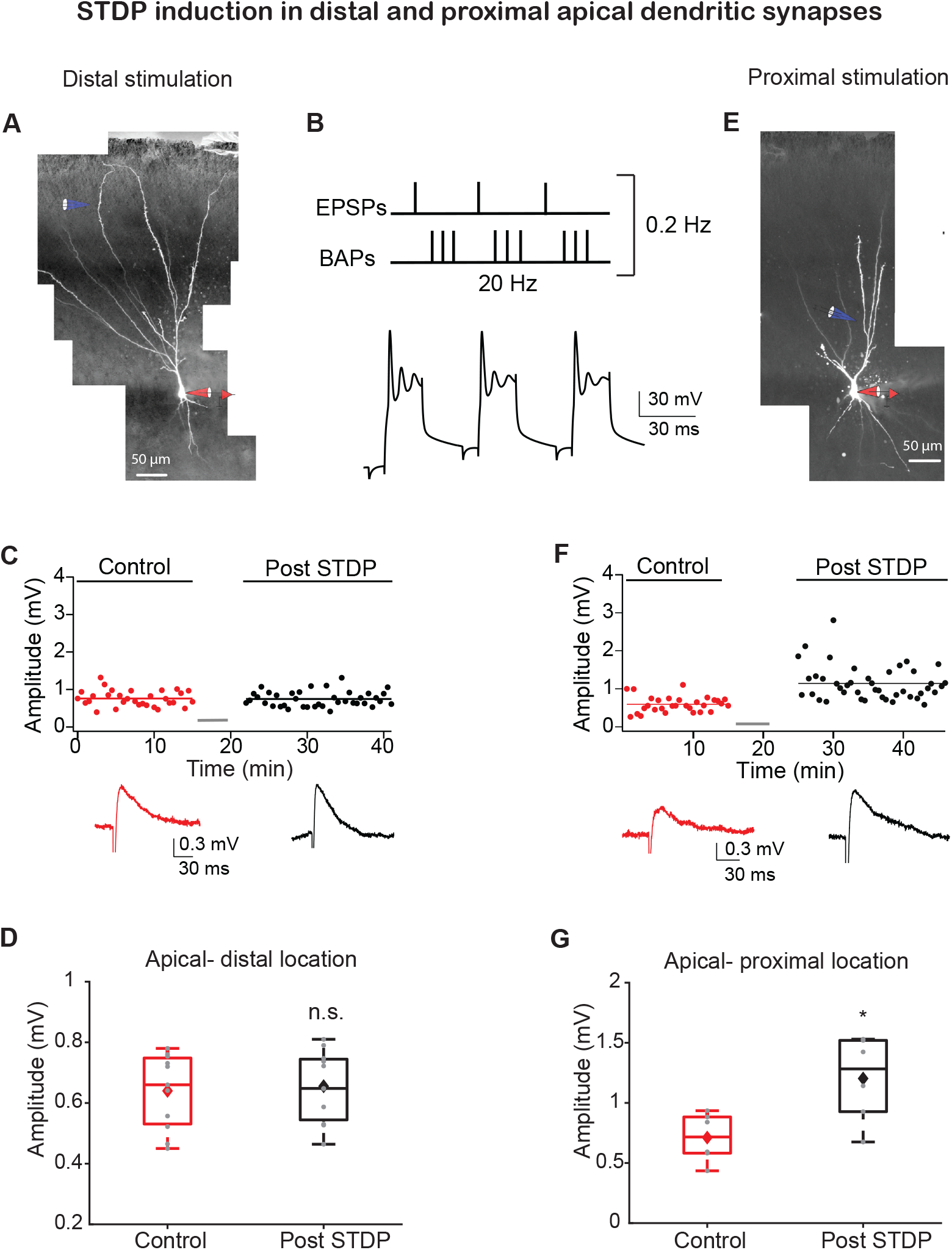
STDP protocol induces LTP in synapses at proximal apical but not distal dendritic locations. **(A)** Experimental setup. A pyramidal neuron from layer IIb was loaded with fluorescent dyes CF-633 (200 µM) and OGB-1 (200 µM) via a somatic patch electrode (red electrode). Focal stimulation was performed using a theta-electrode located nearby a distal apical dendrite in layer Ia (blue electrode; 299 ± 6.29 µm from soma), while recording at the soma. **(B)** STDP induction protocol (top): 1 EPSP followed (8 ms delay) by a burst of 3 BAPs at 150 Hz, repeated 3 times at 20 Hz. Further, this triplet paired EPSP-BAPs was repeated 30 times at 0.2 Hz. (bottom) Example of a voltage response to this induction pairing stimulus. **(C)** Amplitude of single EPSPs is represented over time for control stimulation and after STDP induction protocol at distal apical dendrite (grey bar represents the time of induction stimulus). Control EPSPs were recorded at 0.033 Hz for 10-15 min, prior and after STDP induction protocol. Bottom, traces of average EPSPs for control (red) and post induction (black) from the cell shown in A. **(D)** Box plot showing EPSP amplitudes pre and post STDP induction, for layer Ia inputs. No significant change was observed, post STDP EPSP amplitude was 103.06 ± 2.81% of the control (p = 0.7658; n = 11). **(E)** Same as in A, for activation proximal apical dendrite activation site (blue electrode; 139.33 ± 5.15 µm from soma) at layer Ib. **(F)** Same as in A, for proximal apical dendritic activation site. Control EPSPs were recorded at 0.03 Hz for 10-15 min, prior and after STDP induction protocol. Bottom, traces of average EPSPs for control (red) and post induction (black) from the cell shown in E. **(G)** Box plot showing EPSP amplitudes pre and post STDP induction, for layer Ib inputs. Layer Ib EPSPs were significantly enhanced post induction protocol, 168.93 ± 6.54% of control (p = 0.01387, n = 6). In box plots, the grey dots represent average EPSP of each experiment and the diamond represents mean of the entire set. See also Figure supplement 1.

PCx pyramidal neurons also possess an elaborated basal dendritic arbor (Suzuki and Bekkers, 2006). These dendrites receive IC inputs, mainly feedback inputs from local neurons in PCx and other olfactory areas (Hagiwara et al., 2012; Isaacson, 2010). We focally stimulated inputs innervating basal dendrites (129.25 ± 8.83 µm from the soma, n = 8) using the same STDP protocol as for apical dendrites (Figure 2A). We observed a significant LTP in these basally located synapses, however, with a relatively smaller magnitude (Figure 2B-D; 137.12± 6.11 % of control EPSP; p = 0.0031, n=8) compared to potentiation values in proximal apical dendrites (p= 0.00427), using the same induction protocol.

**Figure 2:**
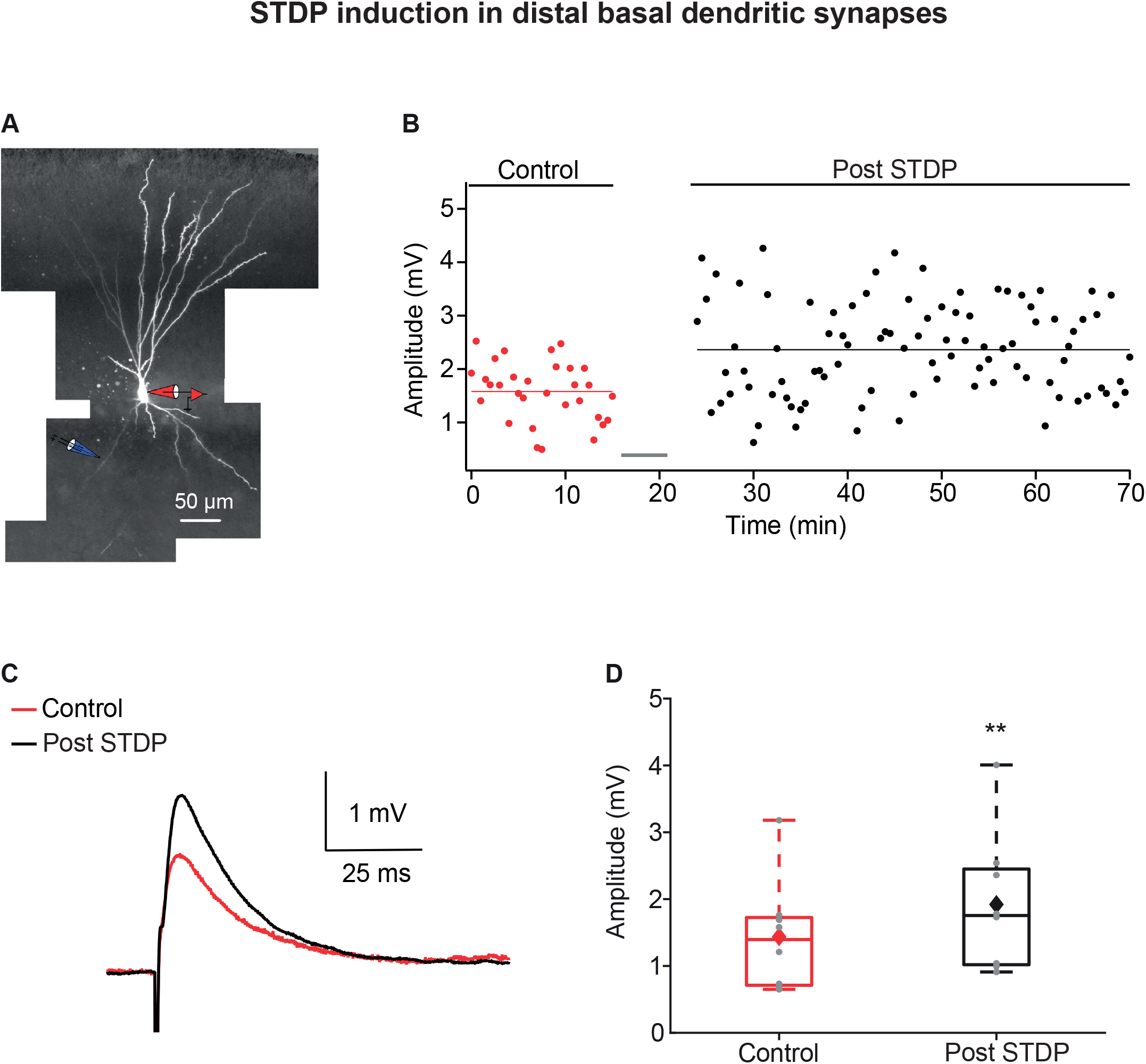
LTP induction via STDP protocol in basal dendrites of PCx pyramidal neurons. **(A)** A layer IIa pyramidal neuron filled with CF633 (200 µM) and OGB-1 (200 µM) via the somatic recording electrode (red electrode). Focal stimulation was performed using a double barrel theta-electrode located nearby a distal basal dendritic site (blue electrode; 129.25 ± 8.83 µm from soma). **(B)** Amplitude of single EPSPs is represented over time for control stimulation and after STDP induction protocol (grey bar represents the time of induction stimulus). Induction protocol, as described in Figure 1B. Bottom, traces of average EPSPs from the cell shown in A, for control (red) and post induction (black). **(C)** Traces of average EPSPs evoked during control (red) and after STDP induction (black). **(D)** Box plot showing EPSP amplitudes pre and post STDP induction during control (red), and post STDP induction (black), for distal basal input stimulation. EPSPs were significantly enhanced post induction protocol (137.12 ± 6.11 % of the control; p = 0.0031, n = 8). The grey dots represent the average EPSP for each cell and the diamond represents mean EPSP of the entire set.

Thus, the STDP burst protocol is efficient in inducing robust LTP in IC inputs both at proximal apical locations and basal synapses but fails altogether to induce LTP at LOT synapses located on distal apical dendrites.

### Local NMDA-spikes induce LTP of LOT inputs in distal dendrites of PCx pyramidal neurons, but fail to potentiate proximal IC inputs

We recently reported the initiation of local NMDA-spikes in distal apical dendrites of PCx pyramidal neurons by activation of LOT synapses (Kumar et al., 2018). Dendritic spikes are excellent candidates for serving as a local postsynaptic signal for induction of long-term synaptic plasticity (Gambino et al., 2014; Golding et al., 2002; Gordon et al., 2006; Lisman and Spruston, 2005, 2010; Major et al., 2013; Polsky et al., 2009; Remy and Spruston, 2007).

To investigate the possible role of NMDA-spikes in inducing LTP of LOT inputs, we recorded from layer II pyramidal neurons, and activated LOT synapses both synaptically and optogenetically. For synaptic stimulation, we focally stimulated LOT synapses impinging on distal layer Ia dendrites (272.75 ± 6.6 µm from the soma) and evoked local NMDA-spikes using a short burst (3-stimuli) at 50 Hz (Figure 3A, 3B). The LTP induction protocol consisted of 2-7 local NMDA-spikes repeated at 4 Hz, mimicking the exploratory sniff cycle (Figure 3C). This local induction protocol was very efficient in inducing robust and strong LTP of EPSPs in the distal LOT synapses terminating in layer Ia (213.98 ± 10.81%; p = 0.0000687; n=26; Figure 3D, 3F). As few as 2 full blown NMDA-spikes at 4 Hz were enough to cause potentiation (Figure 3E), which lasted for the length of the recording (>75 min). To ensure that the distal NMDA-spikes and local EPSPs were indeed evoked primarily by stimulation of LOT inputs, we performed two additional experiments: the first was using the GABA_B_ agonist baclofen (100 µM) which has been shown to preferentially silence IC inputs (Apicella et al., 2010; Franks and Isaacson, 2005; Kumar et al., 2018; Tang and Hasselmo, 1994). Application of Baclofen did not change the resting membrane potential and did not alter the chances of occurrence of NMDA-spikes (Kumar et al., 2018). Bursts of NMDA-spikes delivered at 4 Hz in the presence baclofen also caused potentiation of layer Ia inputs (Figure 4A-D; 236.57 ± 17.51 % of the control EPSP recorded at 0.033 Hz; p = 0.0246; n = 5 cells). In the second set of experiments, we activated LOT inputs optogenetically using ChR2 viral transfection to the bulb (Figure 5). In these experiments EPSPs were generated by optogenetic activation of LOT axons nearby a distal apical dendrite (∼5µm^2^ illumination spot) and NMDA-spikes were generated by either more intense optogenetic activation of LOT inputs or glutamate uncaging (MNI-Glutamate) (Figure 5A-B). Only 3 pairings of opto-EPSPs and NMDA-spikes were sufficient in inducing robust and strong LTP of the LOT inputs (232.45 ± 16.55%; p = 0.000286; n=11; Figure 5A-D).

**Figure 3:**
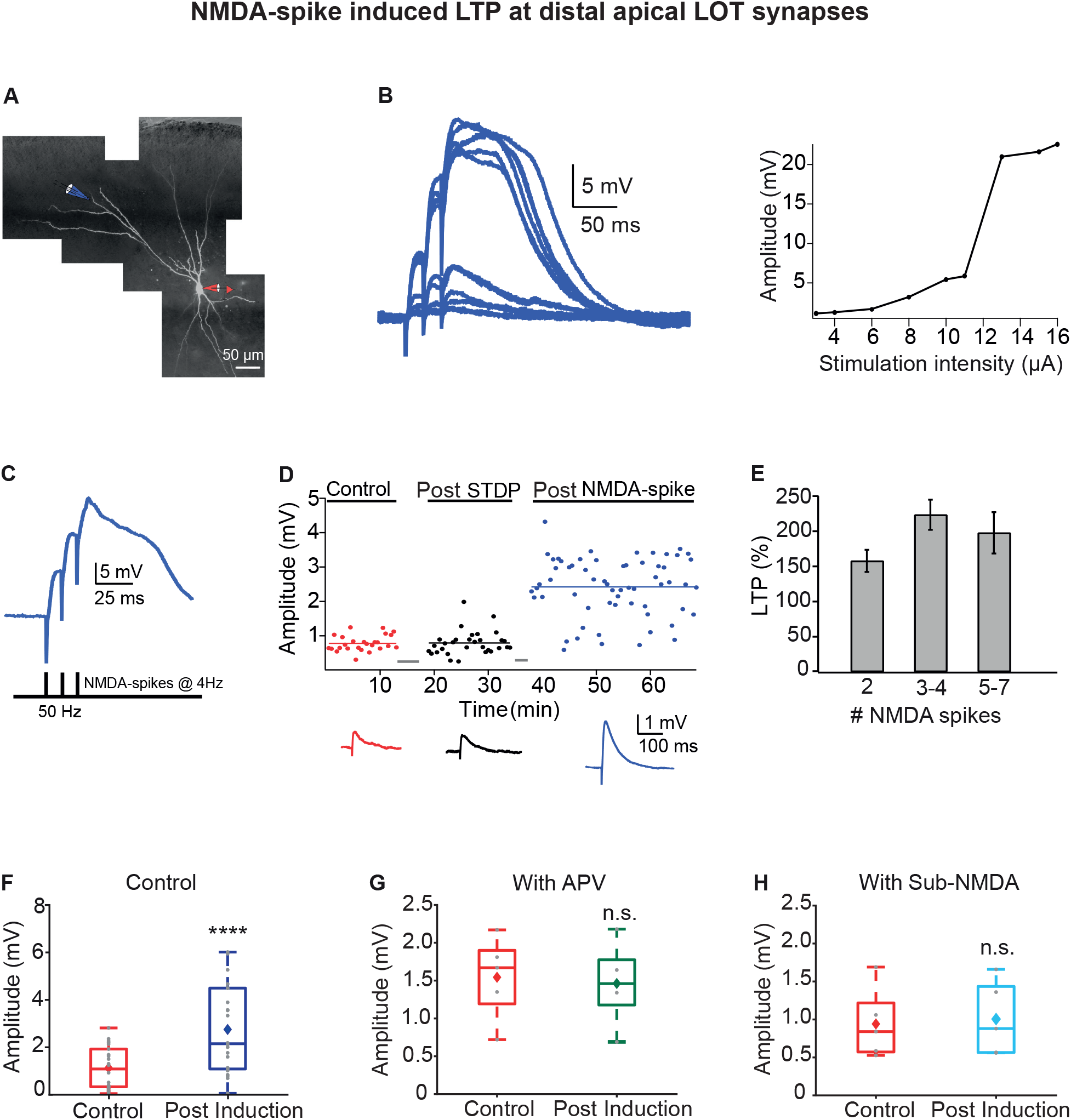
LTP of LOT inputs by NMDA-spikes in PCx pyramidal neurons. **(A)** Fluorescence image of a layer IIa pyramidal neuron filled with CF633 (200 µM) and OGB-1 (200 µM) via the patch recording electrode (red electrode). A focal synaptic stimulating theta-electrode was placed at the distal apical dendrite at layer Ia (blue electrode; 272.75 ± 6.6 µm from the soma). **(B)** Voltage responses evoked by gradually increasing synaptic stimulation intensity (burst of 3 pulses at 50 Hz). Peak voltage response as a function of stimulus intensity showing an all-or-none response (left). **(C)** Schematic of NMDA-spike induction protocol (bottom). NMDA-spikes evoked by 3 pulses at 50 Hz, repeated at 4 Hz for 2-7 times. Upper panel, example voltage response to NMDA-spikes induction protocol. **(D)** Amplitude of single EPSPs is represented over time for control stimulation (red), after STDP induction protocol (black) and after NMDA-spike induction protocol (blue). Grey bars represent the time of induction stimulus. Control EPSPs were recorded at 0.033 Hz for 10-15 min. Potentiation was observed only after the NMDA-spike induction protocol. Bottom, traces of average EPSPs in control (red), post STDP (black) and post NMDA-spike induction (blue), for the cell shown in A. **(E)** Plot of % LTP (relative to control EPSPs) vs number of NMDA-spikes evoked during the induction protocol. All values are insignificant. **(F)** Box plot showing the EPSPs amplitude during control (red) and post NMDA-spike induction protocol (blue). NMDA-spike induction protocol induced large potentiation of the control EPSP (213.98 ± 10.81%; p = 0.0000687; n=26). **(G)** Box plot showing the EPSP amplitude during control NMDA-spike induction protocol (red) and after induction in presence of APV (50 µM; green). No significant change in EPSP was observed when NMDA-spikes were blocked with APV (95.06 ± 4.69 % of control; p = 0.817, n= 5). **(H)** Box plot showing the EPSP amplitude for control NMDA-spike induction protocol (red) and after induction with sub-NMDA EPSPs (teal). No significant change in EPSP amplitudes was observed (106.79 ± 5.69 %, p = 0.8373, n=5). In box plots, the grey dots represent average EPSP for each cell and the diamond represents mean EPSP of the entire set. See also Figure supplement 2.

**Figure 4:**
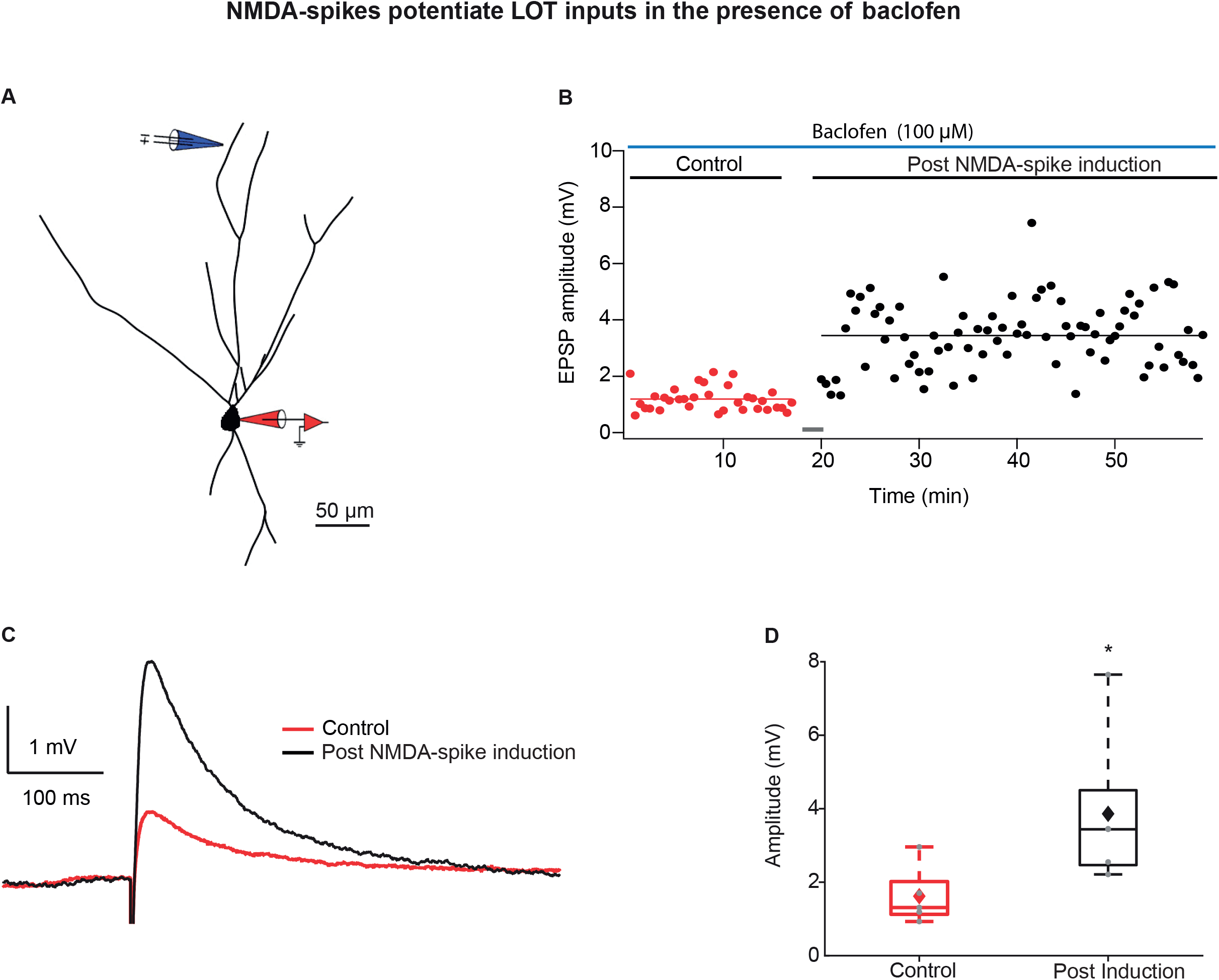
LTP of LOT inputs by NMDA-spikes while blocking IC inputs with baclofen. **(A)** Reconstruction of layer IIa pyramidal neuron filled with CF633 (200 µM) and OGB-1 (200 µM) via the patch recording electrode (red). A focal double-barreled synaptic stimulating electrode was placed at the distal apical dendrite in layer Ia (blue; 287 ± 4.34 µm from soma). **(B)** Amplitude of single EPSPs represented over time for control stimulation and after NMDA-spike induction protocol at distal apical dendrite, in presence of GABA_B_ agonist baclofen (100 µM). Grey bar represents the time of induction stimulus. Control EPSPs were recorded at 0.033 Hz (red), prior and post induction protocol (black). **(C)** Average amplitude of EPSPs during control (red) and post NMDA-spike induction protocol (black). **(D)** Box plot showing EPSPs amplitudes pre (red) and post LTP induction (black), for layer Ia inputs. Layer Ia EPSPs were significantly enhanced post induction protocol in the presence of baclofen, 236.57 ± 17.51% of control EPSP (p = 0.0246, n = 5). The grey dots represent average EPSPs of each experiment and the diamond represents mean of the entire set.

**Figure 5:**
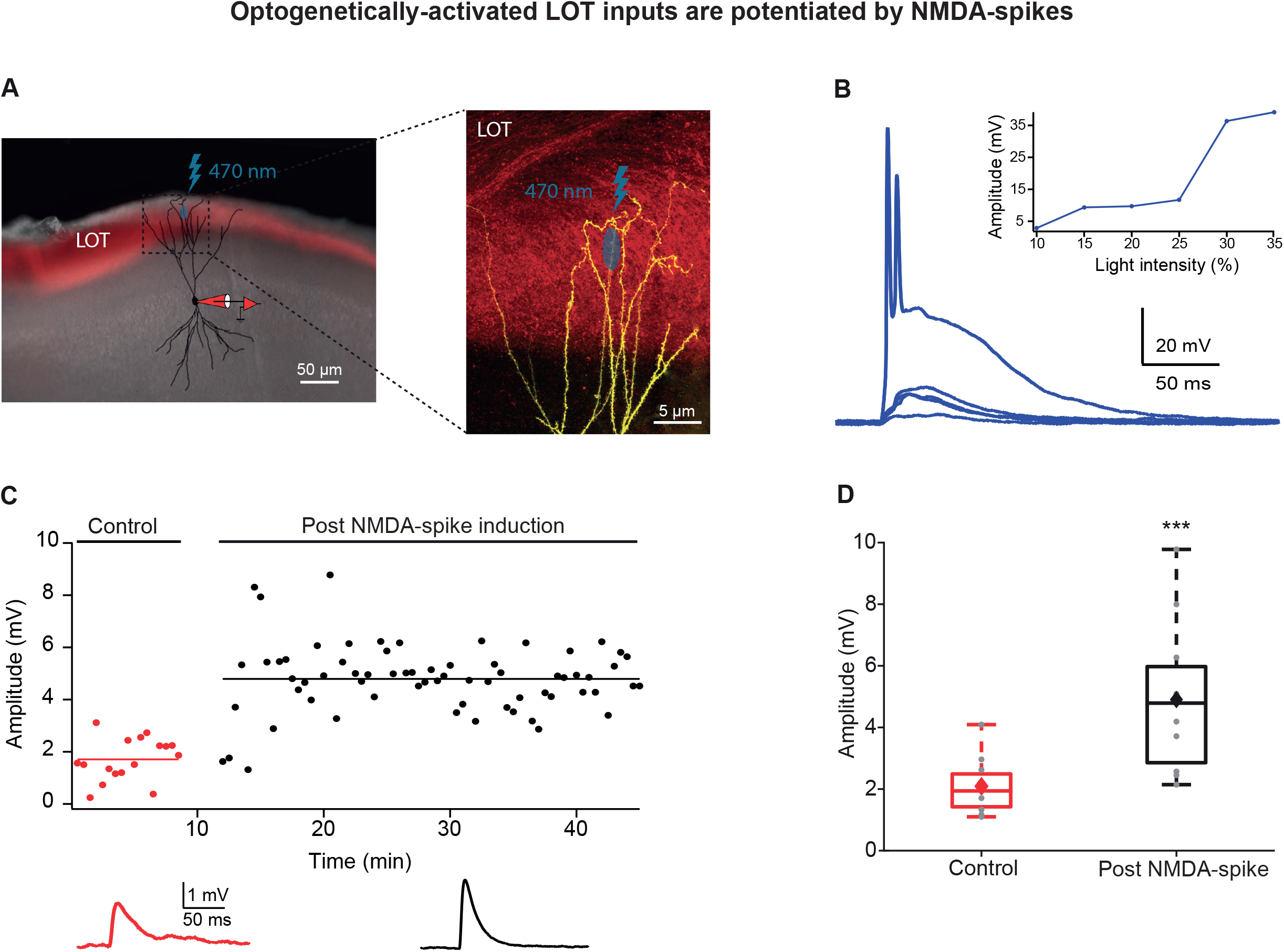
Optogenetically activated LOT inputs were potentiated by NMDA-spikes. **(A)** Experimental setup. Left panel, coronal slice of PCx from a mouse previously injected in OB, with a virus expressing ChR2 (pAAV.CAG.hChR2(H134R)-mCherry.WPRE.SV40) in LOT fibers (red fluorescence). A pyramidal neuron from layer IIb was loaded with CF-633 (200 µM) and OGB-1 (200 µM) via a somatic patch electrode (red electrode) and was reconstructed after the recording session. Right panel, Opto-EPSPs and NMDA-spikes were evoked by light stimulation (LED 470 nm) directed to a small portion of the distal apical dendrite (∼5 µm^2^). **(B)** Voltage responses evoked by gradually increasing ontogenetic light intensity (3 pulses of 5 ms at 50 Hz). Peak voltage responses as a function of % light intensity showing an all-or-none spike response (inset). **(C)** Amplitude of single opto-EPSPs is represented over time for control stimulation and after NMDA-spike induction protocol at distal apical dendrite. Bottom panel, average opto-EPSP in control (red), post NMDA-spike induction (black). **(D)** Box plot showing opto-EPSP amplitudes pre and post NMDA-spike induction, for LOT inputs. Opto-EPSPs were significantly enhanced post NMDA-spike optogenetic induction protocol, 232.45 ± 16.55% of control (p = 0.000286; n=11). The grey dots represent average EPSP of each experiment and the diamond represents mean of the entire set.

We next addressed the question whether NMDA-spikes were critical for potentiation of LOT inputs or whether EPSPs subthreshold to NMDA-spike (sub-NMDA), delivered at 4Hz, can also induce LTP of LOT inputs. EPSPs subthreshold to NMDA-spike initiation delivered with same induction protocol failed to evoke potentiation of the LOT inputs (Figure 3H and Figure supplement 2A-B). The average EPSP amplitude after sub-NMDA-spike induction was 106.79 ± 5.69 % of the control EPSP (p = 0.8373, n= 5 cells). Also, in experiments where NMDA-spikes were blocked by application of the NMDA-R antagonist APV (50 µM), LOT synapses failed to undergo potentiation using the same induction protocol (Figure 3F and Figure supplement 2C-D). The EPSP amplitude post induction was 95.06 ± 4.69 % of control EPSP in the presence of APV (p = 0.817, n= 5).

In contrast to distal LOT synapses, a similar NMDA-spike induction protocol (up to 23 NMDA-spikes at 4Hz) applied at more proximal apical locations by activation of IC synapses (138 ± 3.49 microns from soma) failed to induce potentiation of these IC inputs (Figure 6A-D). Post induction, the proximal IC EPSP amplitude was 99.34 ± 1.8% of the control (p = 0.9886; n = 5).

**Figure 6:**
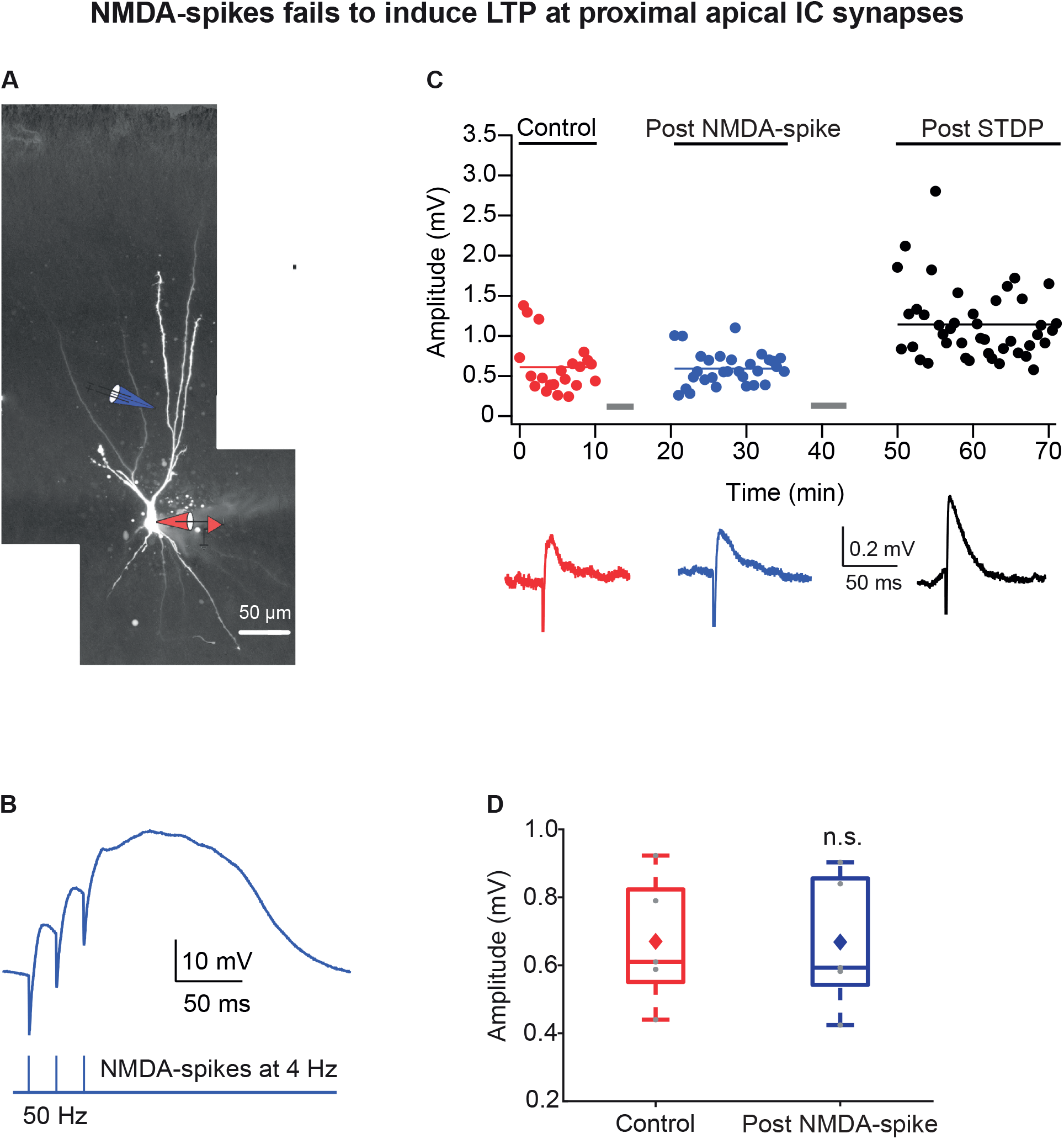
NMDA-spikes fail to induce LTP of IC inputs in layer Ib. **(A)** Fluorescence reconstruction of a layer IIa PCx pyramidal neuron filled with CF633 (200 µM) and OGB-1 (200 µM) via the somatic patch electrode (in red). Focal stimulation was performed using a double barrel theta electrode located nearby a proximal apical site in intracortical layer Ib (in blue; 138 ± 3.49 µm from soma). **(B)** Schematic of NMDA-spike induction protocol: NMDA-spikes evoked by 3 pulses at 50 Hz, repeated at 4 Hz for 5-7 times. Upper panel, an example trace of NMDA-spike delivered at 4Hz during the LTP induction protocol. **(C)** Amplitude of single EPSPs represented over time, during control (red), post NMDA-spike induction (blue) and post STDP induction protocol (black). Grey bars indicate the LTP induction protocols. Lower panel, average EPSP in control (red), post NMDA-spike induction (blue) and post STDP induction (black). **(D)** Box plot showing EPSPs amplitudes during control and post NMDA-spike induction. There was no significant change in EPSP amplitude (99.34 ± 1.8 % of control; p = 0.9886, n=5). The grey dots represent the average EPSP for each cell and the diamond represents mean EPSP of the entire set.

Taken together, these results confirmed that NMDA-spikes are strong mediator for potentiating LOT inputs in layer Ia but were not efficient in potentiating IC inputs at more proximal dendritic locations.

### Local NMDA-spikes can be initiated and induce potentiation in basal dendrites of PCx pyramidal neurons

Basal dendrites in PCx pyramidal neurons are also a major target for IC connections (Haberly, 1985; Isaacson, 2010; Luskin and Price, 1983a, b). Presently, it is unknown whether basal dendrites can generate local NMDA-spikes and in turn whether these spikes can serve to induce plasticity in these dendritic locations.

To test this possibility, we focally stimulated distal basal dendrites using visually positioned theta electrode placed at single basal dendrites with an average distance of 144.83 ± 9.21 µm from the soma (Figure 7A). Similar to apical dendrites, dendritic NMDA-spikes were evoked in basal dendrites of PCx neurons (Figure 7B). The average spike threshold recorded at the soma was 10.2 ± 1.6 mV. The dendritic spike amplitude and area as measured at the soma was 27.2 ± 2.5 mV and 2999.5 ± 434.7 mV*ms respectively (n = 6 cells).

**Figure 7:**
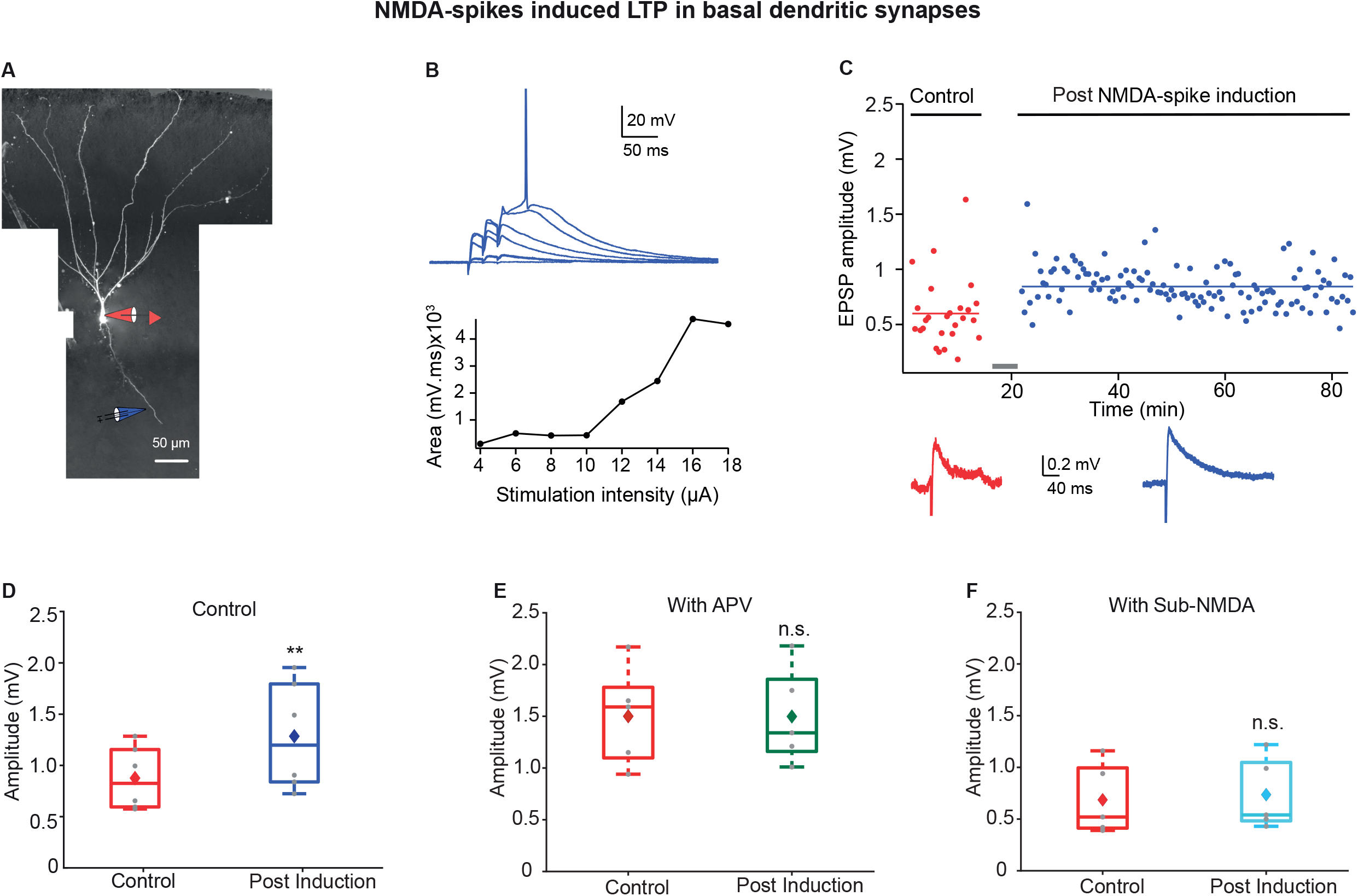
LTP of basal inputs induced by local NMDA-spikes in PCx pyramidal neurons. **(A)** Fluorescence image reconstruction of a layer IIa pyramidal neuron filled with CF633 (200 µM) and OGB-1 (200 µM) via a somatic patch electrode (red electrode). Focal stimulation was performed using a theta electrode placed nearby a distal basal dendrite (blue electrode; 144.83 ± 9.21 µm from soma), while recording at soma. **(B)** Voltage responses evoked by gradually increasing synaptic stimulation consisting of a burst of 3 pulses at 50 Hz. With gradually increasing stimulus intensity, an NMDA-spike was evoked (Top). The peak voltage response is presented as a function of stimulation intensity for the voltage responses shown above (bottom). **(C)** Amplitude of single EPSPs is represented over time for control stimulation and NMDA-spike induction protocol at a basal dendrite. EPSPs were recorded at 0.033 Hz before and after the induction. Down, average EPSPs during control (red) and post NMDA-spike induction paradigm (blue). **(D)** Box plot depicting EPSP amplitudes pre and post-NMDA-spike induction protocol. NMDA-spike induction protocol induced potentiation of the control EPSP, 140.64 ± 4.5 % of control (p = 0.006526, n=6). The grey dots represent the average EPSPs for each cell and the diamond represents mean EPSPs of the entire set. **(E)** Box plot showing EPSP amplitudes pre and post LTP induction protocol in presence of APV. No significant change in EPSP amplitudes was observed, 100.69 ± 4.27 % of control (p = 0.9948, n=5). **(F)** Box plot showing EPSP amplitudes pre and post LTP induction with sub-NMDA EPSPs at 4 Hz. No significant change in EPSP amplitudes was observed, 108.73 ± 3.13 % of control (p = 0.825, n= 5). In the box plots, the grey dots represent average EPSPs of each experiment and the diamond represents the mean of the entire set. See also Figure supplement 3.

Next, we tested if these NMDA-spikes can mediate potentiation of basal dendrite inputs. Using the same NMDA-spike induction protocol used for apical dendrites significantly potentiated the IC synapses on basal dendrites (Figure 7C; 140.64 ± 4.5 % of the control; p = 0.006526, n = 6 cells). However, the degree of potentiation was smaller compared to that of LOT inputs in distal apical dendrites (Figure 7D). EPSPs which were sub-threshold for NMDA-spikes failed to induce significant potentiation of the EPSPs (108.73 ± 3.13 % of the control EPSP, p = 0.8253 n = 5; Figure 7F and Figure supplement 3C-D). The same induction protocol in the presence of NMDA-R blocker APV (50 µM) failed to cause potentiation at these basal synapses (100.69 ± 4.27 % of control, p = 0.9948, n = 5; Figure 7E and Figure supplement 3A-B). These results indicate that local NMDA-spikes are necessary for inducing potentiation of basal inputs.

## Discussion

The piriform cortex was shown to be critical for odor discrimination, recognition and memory and thus, odor memory was attributed to plasticity changes in the PCx (Ghosh et al., 2016; Haberly, 2001; Hasselmo and Barkai, 1995; Saar et al., 2012). However, it is still unclear which synapses and pathways are the target of these plasticity changes. Here we challenge the notion that LOT inputs become “hardwired” after the olfactory critical period (Franks and Isaacson, 2005; Kanter and Haberly, 1993; Poo and Isaacson, 2007), and only IC inputs undergo plasticity changes in adulthood. We show that while LOT inputs contacting distal apical PCx pyramidal neuron dendrites do not undergo significant LTP when paired with somatic output activation using STDP plasticity protocols, they undergo strong and robust LTP when co-activated with other spatially clustered LOT inputs that generate local NMDA-spikes. Only a few (≥2) NMDA-spikes are necessary for full blown LTP induction in these distal LOT synapses. In contrast to distal LOT inputs, IC inputs in more proximal apical dendritic locations, show an inverse picture where locally activated NMDA-spikes fail to potentiate IC inputs, whereas back-propagating action potentials in STDP protocol do successfully produce long term potentiation. Finally, to get a fuller picture regarding location dependent plasticity rules we also investigated IC inputs located in basal dendrites. Interestingly, IC inputs in basal dendrites potentiate by both global STDP protocol and local NMDA-spikes, albeit to a smaller extent compared with synapses in apical dendrites.

### Location dependent plasticity in PCx pyramidal neurons

A key finding described in the literature with respect to the two main PCx pathways, is their differential susceptibility to long term plasticity. While associational IC inputs retain their capability for LTP changes throughout adulthood, LOT inputs were reported to have a critical period during development for long term plasticity changes. Previous work has shown that NMDA-R dependent LTP can be induced robustly in IC synapses both by burst stimulation and by STDP induction protocols (Haberly et al., 1994; Johenning et al., 2009; Kanter et al., 1996; Neville and Haberly, 2004). However, induction of LTP was less consistent with LOT inputs (Haberly et al., 1994; Jung et al., 1990; Neville and Haberly, 2004; Roman et al., 1993) and was either very small and inconsistent (10-15% potentiation) using theta burst protocols (Franks and Isaacson, 2005; Jung et al., 1990; Kanter and Haberly, 1990; Poo and Isaacson, 2007; Roman et al., 1993), or absent using STDP protocols (Johenning et al., 2009).

The failure to induce LTP with STDP protocols in distally located LOT synapses is in line with reports from other types of pyramidal neurons (Gordon et al., 2006; Kampa et al., 2007; Larkum et al., 2009; Letzkus et al., 2006; Nevian et al., 2007; Sandler et al., 2016; Sjostrom and Hausser, 2006; Sjostrom et al., 2008) and is consistent with the severe attenuation of back-propagating action potentials to the distal portions of the apical tree (Bathellier et al., 2009; Johenning et al., 2009; Kumar et al., 2018) and with the high density of A-type potassium channels in the distal apical tree of PCx neurons (Johenning et al., 2009). Here we confirm these findings and show that STDP protocols cannot induce LTP in the distally located LOT synapses but can reliably induce LTP in the more proximally located IC synapses in apical and distal basal trees.

In contrast to the prevailing view, here we show that the local NMDA-spike induction protocol was very efficient in generating robust and powerful LTP of LOT synapses. NMDA-spikes powerfully depolarize the distal locations of the apical tree and are associated with large calcium influx locally (Kumar et al., 2018; Major et al., 2008), and thus serve as a strong postsynaptic signal for induction of plasticity. Our results are consistent with a previous study in CA3 pyramidal neurons showing the necessity of local NMDA-spikes for plasticity in these neurons (Brandalise et al., 2016). Moreover, similar to our findings, previous reports show that dendritic spikes in general can serve as strong and efficient mediators for LTP in CA1 pyramidal neurons (Remy and Spruston, 2007). Importantly, our results are also consistent with in-vivo data showing enhanced strength between the olfactory bulb inputs and the PCx during olfactory learning in adulthood (Chu et al., 2016; Cohen et al., 2008; Cohen et al., 2015; Roman et al., 1993). We assume the reason for the inability of prior studies to induce robust LTP of LOT inputs was related to the fact their induction protocols did not reliably generate local NMDA-spikes in distal apical dendrites (Franks and Isaacson, 2005; Kanter and Haberly, 1993; Poo and Isaacson, 2007).

NMDA-spikes failed to induce LTP in more proximal IC inputs. Thus, IC inputs in adulthood are not merely more prone to undergo plasticity changes compared to LOT synapses as previously suggested (Franks and Isaacson 2005; Poo and Isaacson 2007). It is yet unclear why distally located LOT synapses on apical dendrites undergo strong LTP with NMDA-spikes, while more proximally located IC synapses on apical dendrites cannot. From our findings we cannot determine whether the difference between distal apical LOT and more proximal IC synapses results from different dendritic properties in proximal versus distal apical dendrites, or whether this difference is related to the types of synapses (LOT vs. IC synapses). The location differences cannot be explained simply by the lack of local depolarization or local calcium influx. Rather, other mechanisms need to be considered (for example see Gordon et al. 2006), and further studies are needed to clarify these unknowns. However, regardless of the mechanisms, the computational consequence of our finding indicates that while IC synapses undergo potentiation in response to global activation of the neuron, LOT synapses require co-activation of LOT synapses clustered on the same dendritic segment, and this mechanism in turn allows for storage of different olfactory combinations onto different dendritic branches (Kumar et al., 2018; Poirazi et al. 2003).

### Learning mechanisms in piriform cortex

The anatomical arrangement of the olfactory bulb LOT inputs is such that these inputs terminate broadly throughout the PCx and single pyramidal neurons receive inputs from multiple broadly distributed olfactory glomeruli (Miyamichi et al., 2011; Nagayama et al., 2010; Sosulski et al., 2011). This anatomical arrangement enables a large space of odor combinations to randomly terminate on different neurons and dendritic branches of PCx pyramidal neurons.

A leading theory of odor learning in piriform cortex postulates that an odor representation is generated via strengthening of IC recurrent synapses activated during odor presentation, thus forming odor-specific ensembles in the PCx (Haberly, 2001; Hasselmo and Barkai, 1995; Wilson and Sullivan, 2011). The specificity to odor is enabled by strengthening the connectivity only between neurons that respond to the same set of LOT inputs. Upon exposure to the odor these neurons will fire, and as a result the connectivity between these neurons will be strengthened by associative LTP mechanisms. Upon repeated exposures to the odor, a stronger and more robust activation of this odor-specific neuronal ensemble will take place due to the potentiated IC recurrent connections (Haberly, 2001).

Experimental evidence supports the strengthening of IC inputs by an associative LTP mechanism, where IC and LOT inputs are co-activated (Johenning et al., 2009; Kanter and Haberly, 1993). In support of this idea, plasticity of IC synapses *in vivo* was shown in multiple studies (Ghosh et al., 2015; Saar et al., 2001, 2002; Saar et al., 2012).

Here we propose an additional component to this model, where LOT inputs coding for a specific odor and terminating on the same distal dendritic branch, could be self-potentiated via the NMDA-spike mechanism and a memory trace of the odor will be formed in piriform cortex. Upon reactivation of the same odor, these potentiated odor specific-LOT inputs could recall the odor more easily and further contribute to more efficient recruitment of odor-specific ensembles, ultimately enabling a more sensitive and reliable recall of odors (Weber et al., 2016; Wu and Mel, 2009). Such a mechanism would support the finding *in-vivo* showing potentiation of bulb inputs to PCx during complex olfactory learning (Cohen et al., 2008; Cohen et al., 2015). The plasticity mechanism we propose involving NMDA-spike initiation has physiological feasibility as the number of synapses required for NMDA-spike initiation with in-vivo like firing pattern in PCx pyramidal neurons was estimated to be ∼10-20 clustered synapses in single distal tuft dendrites (Kumar et al., 2018). This number constitutes only a small fraction of the estimated ∼200 LOT synapses contacting distal tuft dendrites of a single neuron (Davison and Ehlers, 2011; Miyamichi et al., 2011).

Association of both potentiated odor information via the LOT, with contextual information from local and from higher order regions such as orbitofrontal cortex and amygdala, can potentially allow the assignment of cognitive and emotional value to odors. Formation of an odor memory trace at the LOT with branch specific dendritic plasticity mechanisms, will further increase the computational and storage capacity of PCx neurons. Moreover, directly strengthening odor representations via such a branch specific plasticity mechanisms of LOT synapses, is expected to enhance the stability of odor memory in the PCx, and to augment the single neuron capacity to represent high-order combinations of odors (Kumar et al., 2018).

## Materials and methods

Key Resources Table.

| **Reagent type (species) or resource** | **Designation** | **Source or reference** | **Identifiers** |
| --- | --- | --- | --- |
| Genetic reagent (M. musculus) | C57BL/6J | Jackson laboratory | RRID: IMSR_JAX:000664 |
| strain, strain background | Wistar (rat)<br>Male/Female | Envigo | RccHan<sup>®</sup>:WIST |
| Recombinant DNA reagent | pAAV.CAG.hChR2(H134R)-mCherry.WPRE.SV40 | Addgene | ID: 100054 |
| Chemical compound | CF-633 Hydrazide | Biotium | Cat.#: 92156 |
| Chemical compound | Oregon Green 488 BAPTA-1 Hexapotassium salt | Invitrogen | Cat.#: 06806 |
| Chemical compound | MNI-Caged-L-glutamate | Tocris Biosciences | Cat.#: 1490/10 |
| Chemical compound | D-AP5 | Tocris Biosciences | Cat.#: 0106/1 |
| Chemical compound | Baclofen | Tocris Biosciences | Cat.#: 0796/10 |
| Software,<br>algorithm | Igor Pro 8 | WaveMetrics | RRID:SCR_000325 |
| Software,<br>algorithm | pClamp 10 | Molecular<br>devices | RRID:SCR_011323 |
| Software,<br>algorithm | Excel | Microsoft | RRID:SCR_016137 |
| Software,<br>algorithm | Adobe Illustrator | Adobe | RRID:SCR_010279 |
| software,<br>algorithm | MATLAB | MathWorks | RRID:SCR_001622 |

### Electrophysiology and Calcium imaging

All animal procedures were done in accordance with guidelines established by NIH on the care and use of animals in research and were confirmed by the Technion Institutional Animal Care and Use Committee.

Acute coronal brain slices 300 µm thick, were prepared from the anterior piriform cortex of Wistar rats (male and female) 4-6 weeks old or mice 7-12 weeks old. The entire brain was removed and placed in ice-cold sucrose solution, maintained under 5° C temperature and saturated with 95% oxygen and 5% CO_2_. The sucrose solution contained (in mM): 2.5 KCl, 1.25 NaH_2_PO_4,_ 25 NaHCO_3_, 7 MgCl_2_, 7 Dextrose, 9 Ascorbic acid, 300 sucrose. The slices were kept in an artificial cerebrospinal fluid (ACSF) at 34-36° C for 30 min recovery, and later kept at room temperature. The ACSF solution contained (in mM): 125 NaCl, 25 Glucose, 3 KCl, 25 NaNCO_3_, 2 CaCl_2_, 1.25 NaH_2_PO_4_, 1 MgCl_2_, pH 7.4.

During experiment, cells were visualized with a confocal scanning microscope equipped with IR illumination and Dodt gradient contrast video microscopy. Whole cell patch clamp recordings were performed on visually identified layer II pyramidal neurons, using an Axon amplifier (Multi-clamp 700A). For patching, glass electrodes (6-9 MΩ) were prepared from thick-walled (0.2 mm) borosilicate glass capillaries (2 mm), using a micropipette puller (P-97, Sutter Instrument). The intracellular pipette solution contained (in mM): 135 K^+^-gluconate, 4 KCl, 4 Mg-ATP, 10 Na_2_-phosphocreatine, 0.3 Na-GTP, 10 HEPES, 0.2 OGB-1, 0.2 CF 633, pH 7.2.

Fluorescence confocal microscopy was performed on an upright BX61WI Olympus microscope equipped with 60X (Olympus 0.9 NA) water-immersion objective. Neurons were filled with calcium-sensitive dye OGB-1 (200 µM; Invitrogen) and CF 633 (200 μM; Biotium) to visualize and collect calcium signals during focal stimulation from the apical and basal dendritic trees. Calcium transients were recorded in a line-scan mode at 500 Hz. The temperature of the slice bath was maintained at 34 °C for the entire duration of the experiment.

### Focal synaptic stimulation

Focal synaptic stimulation at apical and basal dendrites of pyramidal neurons was performed using theta-glass pipette (borosilicate; Hilgenberg), placed in close proximity (5-10 µm) to the desired dendritic segment guided by the fluorescent image of the dendrite and Dodt contrast image of the slice. The theta-stimulation electrodes were filled with CF 633 (0.1 mM; Biotium) diluted with filtered ACSF. Current was delivered through the electrode (short bursts of 3 pulses at 50 Hz) at varying intensities using a stimulus isolator (ISO-Flex; AMPI). The efficacy and location of the stimulation was verified by simultaneous calcium imaging evoked by small EPSPs and their localization to a small segment of the stimulated dendrite.

### Glutamate uncaging

Caged MNI-L-glutamate (Tocris, UK) was delivered locally nearby a dendritic segment in LOT region using pressure ejection (5-10 mbar) from a glass electrode (2-3 µm in diameter) containing 5-10 mM caged glutamate. The electrode was placed 20-30 µm from the dendrite of interest and the caged glutamate was photolyzed using a 1 ms laser pulse (351 nm, Excelsior, Spectra Physics) using point scan mode (Olympus FV1000).

### Drug application

All experiments were performed in the presence of one of the two anion gamma-Aminobutyric acid (GABA_A_) blockers, Bicuculline (1 µM; Sigma) or Gabazine (10 µM; Tocris Bioscience). There were no differences in outcomes observed between these two GABA_A_ blockers. In some experiments, the NMDA-R blocker (APV 50 µM; Tocris Bioscience) was added to the ACSF perfusion solution 20-30 minutes before the start of the recording. GABA_B_ agonist baclofen (100 µM) was added in some experiments to selectively silence IC connections.

### Stereotaxic virus injections

C57B mice 4-7 weeks old, were anaesthetized by isoflurane inhalation (4% for induction and 1.5%– 2% during surgery) and mounted on a stereotaxic frame. Ketoprofen (5 mg/kg) and buprenorphine (0.1 mg/kg) were administered. Labeling of the LOT fibers was performed by unilateral injection of pAAV.CAG.hChR2(H134R)-mCherry.WPRE.SV40 (Addgene) into the OB at coordinates +4.0 mm and +4.5 mm AP from Bregma, +1.0 mm ML at depth 1, 1.5 and 2 mm (20 nL in each depth). The injections were made through the thinned skull using a hydraulic micromanipulator (M0-10 Narishige). Following the surgery, Ketoprofen and Buprenorphine were administered to the animal for 2 consecutive days. The animals were then returned to their home cage for a period of minimum 3 weeks to ensure full recovery from the surgery and expression of the injected virus.

### Long-term plasticity induction protocols

Control EPSPs were acquired at 0.033 Hz before (10-15 minutes) and after the induction protocol. Stability of the recording was ensured by monitoring the resting membrane potential and only experiments in which the membrane potential was not changed more than 3 mV were included. Spike timing dependent plasticity (STDP) was induced by pairing a single EPSP with a burst of post-synaptic back-propagating action potentials (3 BAPs at 150 Hz) evoked by somatic current injection. Triplets of this EPSP-BAPs pairing (at 20 Hz) were repeated 40 times at 5 seconds interval. In some experiments, post-synaptic BAPs were generated by synaptic stimulation of proximal apical dendrites (∼100 microns from soma).

NMDA-spike LTP protocol consisted of 2-23 NMDA-spikes (evoked by a burst of 3 stimuli at 50 Hz) repeated at 4 Hz, the exploratory sniff frequency in rats. Same protocol was repeated with synaptic stimulation which was subthreshold for NMDA-spike initiation (sub-NMDA EPSPs) and in presence of APV (50 µM).

For optogenetic LTP induction protocol, stimulation was performed using a Mightex Polygon micromirror device using a 470 nm blue LED light-source. Light was delivered to the top of the slice through 60X objective (Olympus 0.9 NA). Opto-EPSPs were evoked by 5 ms light pulses delivered every 30 seconds. To evoke an NMDA-spike by optogenetically activating LOT fibers, 3 light pulses of 5 ms each were delivered at 50 Hz.

### Data Analysis and statistical procedure

The sample size was chosen based on standards used in the field with similar experimental paradigms. Importantly most of our experiments involve examining a variable on the same neuron and thus the sources of variability are smaller in these type of experiments.

All electrophysiological data collected were analyzed using Clampfit 10.3 (Axon instruments), Igor Pro software (5.01, Wave metrics), Microsoft office 365 and an in-house MATLAB software. All data is presented as mean ± SEM. Two-tailed paired Student’s t-test was used for testing statistical significance of the data. All neurons in which the recordings deteriorated, as measured by the series resistance of the electrode, were excluded from the averages.

## Data availability

All relevant data are included in the paper and/or its supplementary information files. Additional data will be made available from the corresponding author upon reasonable request.

## Acknowledgements

We thank Y. Schiller for helpful discussions throughout the project and helpful comments on the manuscript and B. Mel for helpful comments on the manuscript. We thank Irena Reiter for excellent technical assistance. This study was supported by Israeli Science Foundation (J.S.), German Israeli Foundation (J.S.) and Prince funds (J.S.).

## Author Contributions

Conceptualization, J.S. E.B.; Methodology, A.K., J.S.; Formal Analysis, A.K., J.S.; Investigation, A.K., J.S.; Data Curation, J.S., A.K.; Writing, J.S., A.K., E.B.; Visualization, A.K. and J.S.; Funding Acquisition, J.S.

## Supplementary information

**Figure supplement 1:**
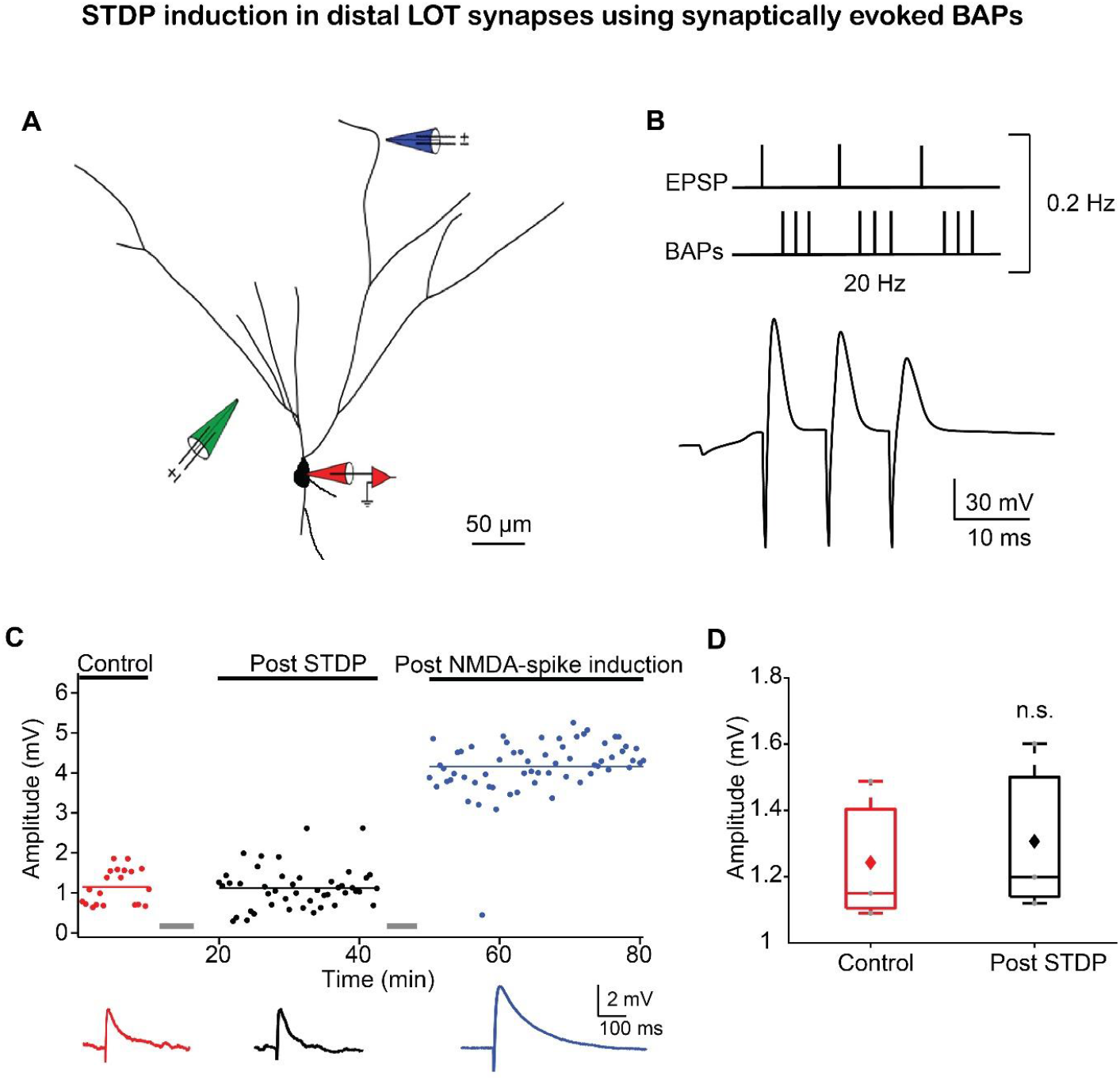
Pairing synaptically induced EPSPs and BAPs also failed to induce LTP of LOT inputs. Related to Figure 1. **(A)** Reconstruction of a layer IIb pyramidal neuron filled with fluorescent dyes CF-633 (200 µM) and OGB-1 (200 µM) via somatic recording electrode (red electrode) and focally stimulated in apical distal dendrite using a theta electrode (blue electrode; 286 ± 33.86 µm from soma). Synaptic EPSPs were evoked by a theta electrode placed in layer IIa, close to the soma (green electrode; 95 ± 7.64 µm from soma). **(B)** The induction protocol was same as described in Figure 1B, except that the BAPs were evoked by synaptic stimulation close to the soma. Bottom, example trace of STDP induction paradigm. **(C)** Amplitude of single EPSPs represented over time for control (red), post STDP induction (black) and post NMDA-spike induction protocol (blue). Pairing EPSPs and synaptically induced BAPs failed to induce LTP. Subsequent delivery of NMDA-spikes at 4 Hz potentiated layer Ia inputs. Bottom panel, average EPSPs for control (red), post STDP (black) and post NMDA-spike induction protocol (blue). **(D)** Box plot showing EPSP amplitudes pre and post STDP, for layer Ia inputs, when BAPs were initiated using synaptic stimulation close to the soma. Layer Ia EPSPs did not change significantly post STDP induction (104.99 ± 3.86% of control EPSP; p = 0.7577, n = 3). The grey dots represent average EPSPs of each experiment and the diamond represents mean of the entire set.

**Figure supplement 2:**
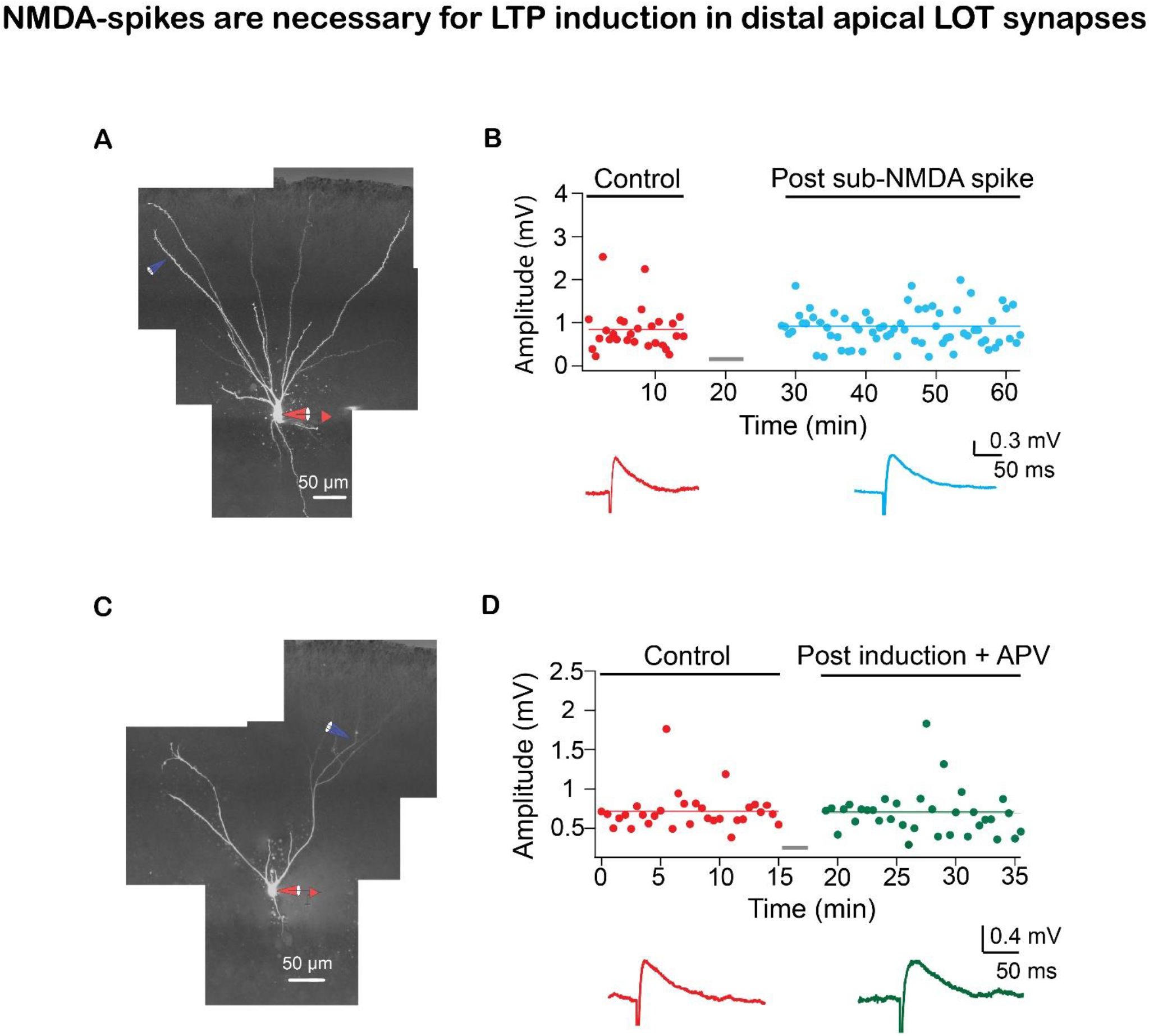
Example experiments showing NMDA-spikes are necessary for LTP of LOT synapses. Related to Figure 3. **(A)** Reconstruction of a layer IIa pyramidal neuron filled with CF633 (200 µM) and OGB-1 (200 µM) via the somatic recording electrode (red) and focally stimulated using a theta electrode in distal apical dendrite at layer Ia (blue; 298 ± 14.38 µm from soma). **(B)** Amplitude of single EPSPs is represented over time for control stimulation (red) and after EPSPs subthreshold to NMDA-spike induction protocol (teal). Same induction protocol as in Figure 3, except stimulation was just subthreshold to NMDA-spikes initiation. Bottom panel, averaged EPSPs evoked during control (red) and post induction with sub-NMDA EPSPs (teal). **(C)** Reconstruction of a layer IIa pyramidal neuron filled with CF633 (200 µM) and OGB-1 (200 µM) via the somatic recording electrode (red) and focally stimulated using a theta electrode in distal apical dendrite at layer Ia (blue; 275.6 ± 11.55 µm from soma). **(D)** Amplitude of single EPSPs is represented over time for control stimulation (red), after NMDA-spike induction protocol in the presence of APV (50 µM; green). Same induction protocol as in Figure 3. Bottom panel, averaged EPSPs evoked during control (red) and post induction in the presence of APV (green).

**Figure supplement 3:**
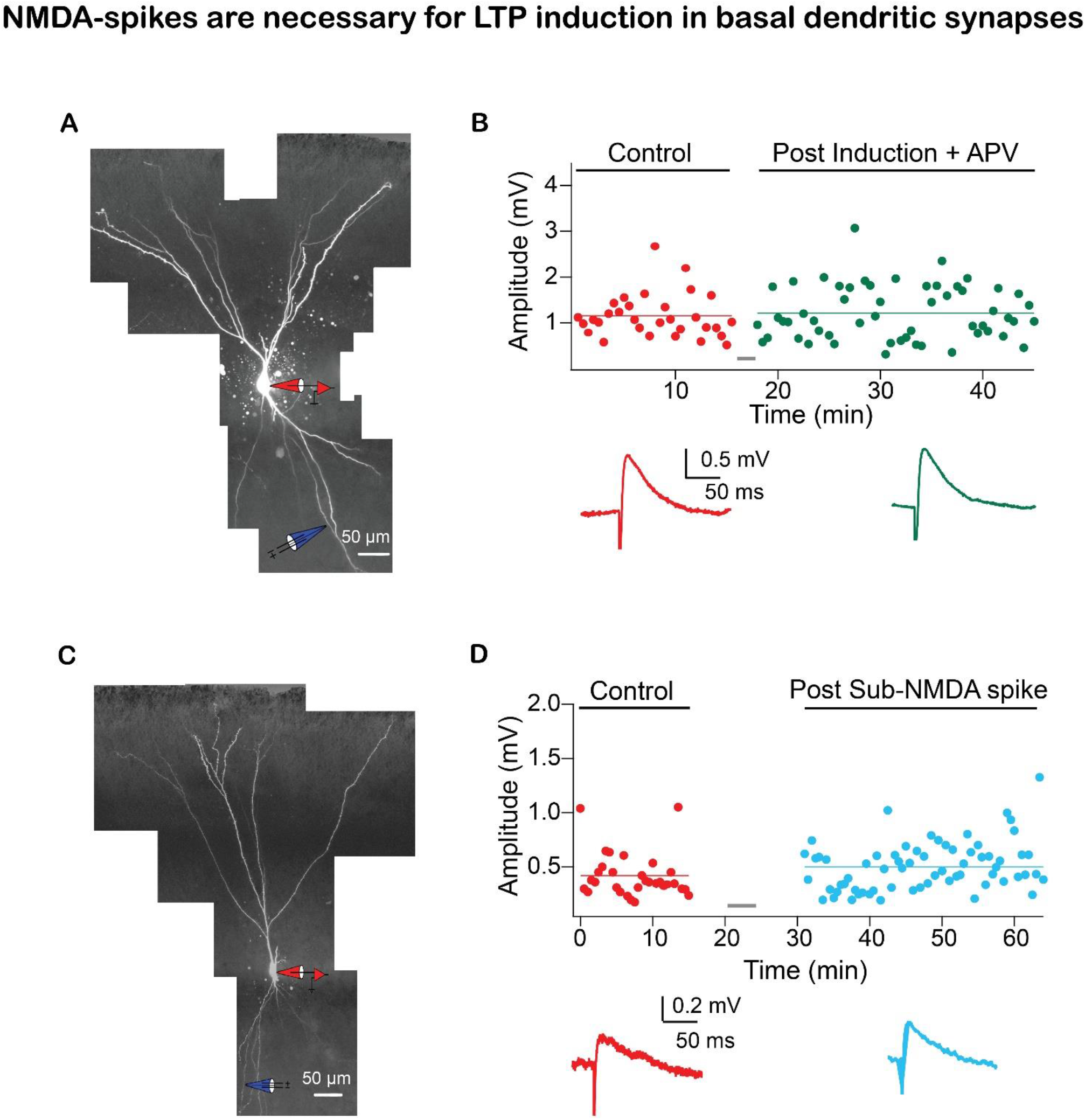
Example experiments showing NMDA-spikes are necessary for LTP of IC synapses in basal dendrites. Related to Figure 7. **(A)** Reconstruction of a layer IIa pyramidal neuron filled with fluorescence dyes CF633 (200 µM) and OGB-1 (200 µM) via somatic patch electrode (red electrode) and focally stimulated using a theta electrode in distal apical dendrite at layer Ia (blue; 174.4 ± 11.82 µm from soma). **(B)** Amplitude of single EPSPs is represented over time for control (red) and post EPSPs NMDA-spike induction protocol (green) in presence of APV (50 µM). No significant change in EPSP amplitude was observed. Grey bar represents the time of induction stimulus. Bottom panel, traces of average EPSPs for control (red) and post induction (green). **(C)** Reconstruction of a layer IIa pyramidal neuron filled with fluorescence dyes CF633 (200 µM) and OGB-1 (200 µM) via somatic patch electrode (red) and focally stimulated at the distal basal dendrite using a theta electrode (blue; 154.8 ± 12.52 µm from soma). **(D)** Amplitude of single EPSPs represented over time, for control (red) and post sub-NMDA spike induction (teal). Grey bar indicates the time of induction stimulus. Bottom panel, average EPSP amplitude during control (red) and post sub-NMDA induction protocol (teal).

## References

Apicella, A., Yuan, Q., Scanziani, M., and Isaacson, J.S. (2010). Pyramidal cells in piriform cortex receive convergent input from distinct olfactory bulb glomeruli. J Neurosci 30, 14255–14260.

Bathellier, B., Margrie, T.W., and Larkum, M.E. (2009). Properties of piriform cortex pyramidal cell dendrites: implications for olfactory circuit design. J Neurosci 29, 12641–12652.

Bekkers, J.M., and Suzuki, N. (2013). Neurons and circuits for odor processing in the piriform cortex. Trends Neurosci 36, 429–438.

Bliss, T.V., and Collingridge, G.L. (1993). A synaptic model of memory: long-term potentiation in the hippocampus. Nature 361, 31–39.

Brandalise, F., Carta, S., Helmchen, F., Lisman, J., and Gerber, U. (2016). Dendritic NMDA spikes are necessary for timing-dependent associative LTP in CA3 pyramidal cells. Nat Commun 7, 13480.

Chu, M.W., Li, W.L., and Komiyama, T. (2016). Balancing the Robustness and Efficiency of Odor Representations during Learning. Neuron 92, 174–186.

Cohen, Y., Reuveni, I., Barkai, E., and Maroun, M. (2008). Olfactory learning-induced long-lasting enhancement of descending and ascending synaptic transmission to the piriform cortex. J Neurosci 28, 6664–6669.

Cohen, Y., Wilson, D.A., and Barkai, E. (2015). Differential modifications of synaptic weights during odor rule learning: dynamics of interaction between the piriform cortex with lower and higher brain areas. Cereb Cortex 25, 180–191.

Davison, I.G., and Ehlers, M.D. (2011). Neural circuit mechanisms for pattern detection and feature combination in olfactory cortex. Neuron 70, 82–94.

Feldman, D.E. (2012). The spike-timing dependence of plasticity. Neuron 75, 556–571.

Franks, K.M., and Isaacson, J.S. (2005). Synapse-specific downregulation of NMDA receptors by early experience: a critical period for plasticity of sensory input to olfactory cortex. Neuron 47, 101–114.

Gambino, F., Pages, S., Kehayas, V., Baptista, D., Tatti, R., Carleton, A., and Holtmaat, A. (2014). Sensory-evoked LTP driven by dendritic plateau potentials in vivo. Nature 515, 116–119.

Ghosh, S., Reuveni, I., Barkai, E., and Lamprecht, R. (2016). Calcium/calmodulin-dependent kinase II activity is required for maintaining learning-induced enhancement of alpha-amino-3-hydroxy-5-methyl-4-isoxazolepropionic acid receptor-mediated synaptic excitation. Journal of neurochemistry 136, 1168–1176.

Ghosh, S., Reuveni, I., Lamprecht, R., and Barkai, E. (2015). Persistent CaMKII activation mediates learning-induced long-lasting enhancement of synaptic inhibition. J Neurosci 35, 128–139.

Golding, N.L., Staff, N.P., and Spruston, N. (2002). Dendritic spikes as a mechanism for cooperative long-term potentiation. Nature 418, 326–331.

Gordon, U., Polsky, A., and Schiller, J. (2006). Plasticity compartments in basal dendrites of neocortical pyramidal neurons. J Neurosci 26, 12717–12726.

Haberly, L., Ketchum, K., and Kanter, E. (1994). LTP in piriform cortex: characterization and functional speculations (Long-term Potentiation: A Debate of Current Issues. MIT Press, Cambridge).

Haberly, L.B. (1985). Neuronal circuitry in olfactory cortex: anatomy and functional implications. Chemical senses.

Haberly, L.B. (2001). Parallel-distributed processing in olfactory cortex: new insights from morphological and physiological analysis of neuronal circuitry. Chemical senses 26, 551–576.

Haberly, L.B., and Price, J.L. (1977). The axonal projection patterns of the mitral and tufted cells of the olfactory bulb in the rat. Brain Res 129, 152–157.

Hagiwara, A., Pal, S.K., Sato, T.F., Wienisch, M., and Murthy, V.N. (2012). Optophysiological analysis of associational circuits in the olfactory cortex. Front Neural Circuits 6, 18.

Hasselmo, M.E., and Barkai, E. (1995). Cholinergic modulation of activity-dependent synaptic plasticity in the piriform cortex and associative memory function in a network biophysical simulation. J Neurosci 15, 6592–6604.

Isaacson, J.S. (2010). Odor representations in mammalian cortical circuits. Curr Opin Neurobiol 20, 328–331.

Johenning, F.W., Beed, P.S., Trimbuch, T., Bendels, M.H., Winterer, J., and Schmitz, D. (2009). Dendritic compartment and neuronal output mode determine pathway-specific long-term potentiation in the piriform cortex. J Neurosci 29, 13649–13661.

Jung, M.W., Larson, J., and Lynch, G. (1990). Long-term potentiation of monosynaptic EPSPs in rat piriform cortex in vitro. Synapse 6, 279–283.

Kampa, B.M., Letzkus, J.J., and Stuart, G.J. (2007). Dendritic mechanisms controlling spike-timing-dependent synaptic plasticity. Trends Neurosci 30, 456–463.

Kanter, E.D., and Haberly, L.B. (1990). NMDA-dependent induction of long-term potentiation in afferent and association fiber systems of piriform cortex in vitro. Brain Res 525, 175–179.

Kanter, E.D., and Haberly, L.B. (1993). Associative long-term potentiation in piriform cortex slices requires GABAA blockade. J Neurosci 13, 2477–2482.

Kanter, E.D., Kapur, A., and Haberly, L.B. (1996). A dendritic GABAA-mediated IPSP regulates facilitation of NMDA-mediated responses to burst stimulation of afferent fibers in piriform cortex. J Neurosci 16, 307–312.

Kumar, A., Schiff, O., Barkai, E., Mel, B.W., Poleg-Polsky, A., and Schiller, J. (2018). NMDA spikes mediate amplification of inputs in the rat piriform cortex. eLife 7.

Larkum, M.E., Nevian, T., Sandler, M., Polsky, A., and Schiller, J. (2009). Synaptic integration in tuft dendrites of layer 5 pyramidal neurons: a new unifying principle. Science 325, 756–760.

Lebel, D., Grossman, Y., and Barkai, E. (2001). Olfactory learning modifies predisposition for long-term potentiation and long-term depression induction in the rat piriform (olfactory) cortex. Cereb Cortex 11, 485–489.

Letzkus, J.J., Kampa, B.M., and Stuart, G.J. (2006). Learning rules for spike timing-dependent plasticity depend on dendritic synapse location. J Neurosci 26, 10420–10429.

Letzkus, J.J., Kampa, B.M., and Stuart, G.J. (2007). Does spike timing-dependent synaptic plasticity underlie memory formation? Clin Exp Pharmacol Physiol 34, 1070–1076.

Lisman, J., and Spruston, N. (2005). Postsynaptic depolarization requirements for LTP and LTD: a critique of spike timing-dependent plasticity. Nat Neurosci 8, 839–841.

Lisman, J., and Spruston, N. (2010). Questions about STDP as a General Model of Synaptic Plasticity. Front Synaptic Neurosci 2, 140.

Luskin, M.B., and Price, J.L. (1983a). The laminar distribution of intracortical fibers originating in the olfactory cortex of the rat. J Comp Neurol 216, 292–302.

Luskin, M.B., and Price, J.L. (1983b). The topographic organization of associational fibers of the olfactory system in the rat, including centrifugal fibers to the olfactory bulb. J Comp Neurol 216, 264–291.

Major, G., Larkum, M.E., and Schiller, J. (2013). Active properties of neocortical pyramidal neuron dendrites. Annu Rev Neurosci 36, 1–24.

Major, G., Polsky, A., Denk, W., Schiller, J., and Tank, D.W. (2008). Spatiotemporally graded NMDA spike/plateau potentials in basal dendrites of neocortical pyramidal neurons. Journal of Neurophysiology 99, 2584–2601.

Miyamichi, K., Amat, F., Moussavi, F., Wang, C., Wickersham, I., Wall, N.R., Taniguchi, H., Tasic, B., Huang, Z.J., He, Z., et al. (2011). Cortical representations of olfactory input by trans-synaptic tracing. Nature 472, 191–196.

Nagayama, S., Enerva, A., Fletcher, M.L., Masurkar, A.V., Igarashi, K.M., Mori, K., and Chen, W.R. (2010). Differential axonal projection of mitral and tufted cells in the mouse main olfactory system. Front Neural Circuits 4.

Nevian, T., Larkum, M.E., Polsky, A., and Schiller, J. (2007). Properties of basal dendrites of layer 5 pyramidal neurons: a direct patch-clamp recording study. Nat Neurosci 10, 206–214.

Neville, K.R., and Haberly, L.B. (2004). Olfactory cortex. The synaptic organization of the brain 5, 415–454.

Patneau, D.K., and Stripling, J.S. (1992). Functional correlates of selective long-term potentiation in the olfactory cortex and olfactory bulb. Brain Res 585, 219–228.

Polsky, A., Mel, B., and Schiller, J. (2009). Encoding and decoding bursts by NMDA spikes in basal dendrites of layer 5 pyramidal neurons. J Neurosci 29, 11891–11903.

Poo, C., and Isaacson, J.S. (2007). An early critical period for long-term plasticity and structural modification of sensory synapses in olfactory cortex. J Neurosci 27, 7553–7558.

Remy, S., and Spruston, N. (2007). Dendritic spikes induce single-burst long-term potentiation. Proc Natl Acad Sci U S A 104, 17192–17197.

Roman, F.S., Chaillan, F.A., and Soumireu-Mourat, B. (1993). Long-term potentiation in rat piriform cortex following discrimination learning. Brain Res 601, 265–272.

Saar, D., and Barkai, E. (2003). Long-term modifications in intrinsic neuronal properties and rule learning in rats. Eur J Neurosci 17, 2727–2734.

Saar, D., and Barkai, E. (2009). Long-lasting maintenance of learning-induced enhanced neuronal excitability: mechanisms and functional significance. Mol Neurobiol 39, 171–177.

Saar, D., Grossman, Y., and Barkai, E. (1998). Reduced after-hyperpolarization in rat piriform cortex pyramidal neurons is associated with increased learning capability during operant conditioning. Eur J Neurosci 10, 1518–1523.

Saar, D., Grossman, Y., and Barkai, E. (2001). Long-lasting cholinergic modulation underlies rule learning in rats. J Neurosci 21, 1385–1392.

Saar, D., Grossman, Y., and Barkai, E. (2002). Learning-induced enhancement of postsynaptic potentials in pyramidal neurons. J Neurophysiol 87, 2358–2363.

Saar, D., Reuveni, I., and Barkai, E. (2012). Mechanisms underlying rule learning-induced enhancement of excitatory and inhibitory synaptic transmission. J Neurophysiol 107, 1222–1229.

Sandler, M., Shulman, Y., and Schiller, J. (2016). A Novel Form of Local Plasticity in Tuft Dendrites of Neocortical Somatosensory Layer 5 Pyramidal Neurons. Neuron 90, 1028–1042.

Sjostrom, P.J., and Hausser, M. (2006). A cooperative switch determines the sign of synaptic plasticity in distal dendrites of neocortical pyramidal neurons. Neuron 51, 227–238.

Sjostrom, P.J., Rancz, E.A., Roth, A., and Hausser, M. (2008). Dendritic excitability and synaptic plasticity. Physiol Rev 88, 769–840.

Sjostrom, P.J., Turrigiano, G.G., and Nelson, S.B. (2007). Multiple forms of long-term plasticity at unitary neocortical layer 5 synapses. Neuropharmacology 52, 176–184.

Sosulski, D.L., Bloom, M.L., Cutforth, T., Axel, R., and Datta, S.R. (2011). Distinct representations of olfactory information in different cortical centres. Nature 472, 213–216.

Suzuki, N., and Bekkers, J.M. (2006). Neural coding by two classes of principal cells in the mouse piriform cortex. J Neurosci 26, 11938–11947.

Suzuki, N., and Bekkers, J.M. (2011). Two layers of synaptic processing by principal neurons in piriform cortex. J Neurosci 31, 2156–2166.

Tang, A.C., and Hasselmo, M.E. (1994). Selective suppression of intrinsic but not afferent fiber synaptic transmission by baclofen in the piriform (olfactory) cortex. Brain Res 659, 75–81.

Weber, J.P., Andrasfalvy, B.K., Polito, M., Mago, A., Ujfalussy, B.B., and Makara, J.K. (2016). Location-dependent synaptic plasticity rules by dendritic spine cooperativity. Nat Commun 7, 11380.

Wilson, D.A. (1998). Habituation of odor responses in the rat anterior piriform cortex. J Neurophysiol 79, 1425–1440.

Wilson, D.A., and Sullivan, R.M. (2011). Cortical processing of odor objects. Neuron 72, 506–519.

Wu, X.E., and Mel, B.W. (2009). Capacity-enhancing synaptic learning rules in a medial temporal lobe online learning model. Neuron 62, 31–41.

